# Fungi form interkingdom microbial communities in the primordial human gut that develop with gestational age

**DOI:** 10.1101/621235

**Authors:** Kent A. Willis, John H. Purvis, Erin D. Myers, Michael M. Aziz, Ibrahim Karabayir, Charles K. Gomes, Brian M. Peters, Oguz Akbilgic, Ajay J. Talati, Joseph F. Pierre

**Author notes:** Correspondence and request for materials should be addressed to J.F.P.

## Abstract

Fungal and bacterial commensal organisms play a complex role in the health of the human host. Expansion of commensal ecology after birth is a critical period in human immune development. However, the initial fungal colonization of the primordial gut remains undescribed. To investigate primordial fungal ecology, we performed amplicon sequencing and culture-based techniques of first-pass meconium, which forms in the fetal intestine prior to birth, from a prospective observational cohort of term-born and preterm newborns. Here, we describe fungal ecologies in the primordial gut that develop complexity with advancing gestational age at birth. Our findings suggest homeostasis of fungal commensals may represent an important aspect of human biology present even before birth. Unlike bacterial communities which gradually develop complexity, the domination of the fungal communities of some preterm infants by Saccromycetes, specifically *Candida*, may suggest a pathologic association with preterm birth.

## Introduction

Human health is intimately entwined with the commensal microorganisms that constitutionally colonize our barrier surfaces, such as the intestinal tract. The human gut microbiome is host to a complex interkingdom ecology of bacteria, fungi, archaea, protozoa and viruses. While recent research has led to a drastic expansion in our understanding of bacterial communities in the gut^1-4^, little is known about the role of fungal organisms, collectively the mycobiome, particularly in relation to their impact on the population dynamics of early-life colonization.

The tightly choreographed maturation of commensal microbial communities is crucial to mammalian immune development^5^. Fungal organisms can form a crucial nidus for initial colonization^6^ and may play a vital role in facilitating early colonization of the neonatal gut. Disruption of this process could produce pathogenic fungal overgrowth or invasive disease. Indeed, this mechanism may play a role in the development of asthma and other metabolic and immunological diseases^7^. The intestinal mycobiota likely confers unique physiologic effects to the host, in part by processing nutrients and vitamins either separately or in conjunction with the microbiome. However, characterization of the functional impact of early mycobiome is impaired by limited knowledge of the normal development of these communities. To understand the functional impact of the early-life mycobiome on human health, it is necessary to determine if the human primordial gut hosts fungal communities, if these communities are established before or after birth, and if the presence of fungi in the early gut impacts neonatal physiology.

Advances in next-generation sequencing techniques have pushed back the timing of initial commensal colonization^3^ which was initially thought to occur at birth^8,9^. Despite these advances, interkingdom interactions within microbial communities and the influence of environmental factors on the dynamics of microbial ecosystems remains poorly understood. Due to delayed progress in sequencing and characterizing fungal communities^10^, the appreciation of commensal fungal ecology and biomedical impact of fungi lags behind the bacterial microbiome. It appears that during the first several years of life, early fungal communities, like their bacterial counterparts, arise from maternal communities, but unlike their bacterial counterparts, remain at a relatively low and consistent diversity^11,12^. The relative consistency of early mycobiome makes understanding initial colonization important to unravel both the long-term biomedical impact of fungal communities and to explain their divergence from bacterial communities. While we are beginning to understand the development of the mycobiome during early-life, the timing of initial fungal colonization remains undescribed.

To characterize the interkingdom ecologies of the fetal microbiome and understand when fungal colonization first occurs, we performed a molecular and culture-based survey of first-pass meconium collected from an observational cohort of preterm and term-born human newborns. In this study, we characterize fungal communities in first-pass meconium within hours of birth and explore the influence of perinatal factors on the foundational ecology of the human gut, laying the groundwork for understanding the functional impact of fungal communities on the host. We support our observations by examining the fetal and neonatal mouse mycobiome. We describe complex bacterial and fungal commensal communities that develop in abundance and diversity with length of gestation. Our results suggest that the human fetus is naturally exposed an increasing variety of microbial DNA as gestation progresses. However, the increased prevalence of fungal order Saccharomyces, especially the genus *Candida*, in preterm infants may also suggest a pathologic association with preterm birth.

## Results

### Study design and cohort characteristics

To characterize the fungal ecology of the primordial human gut, we collected the first-pass meconium from a cohort of very low birthweight preterm (<1500 g, gestational age 23-32 weeks, n = 54) and term-born (gestational age 37-41 weeks, n = 36) newborns and characterized the microbial communities using both culture-independent and culture-dependent techniques (Fig. 1). Of these 90 infants prospectively enrolled from a Level IIIB Neonatal Intensive Care Unit and Well Baby Nursery at single academic medical center, 19 infants (31.5% of preterm and 5.6% of term-born infants, χ^2^ *p =*0.0033) did not produce a meconium stool that could be collected during the first 48 h of life (Fig. 1a). To differentiate the effects of perinatal factors on microbial ecology, we prospectively collected clinical data describing perinatal exposures (Fig. 1b). In addition to lower gestational age, infants born preterm were more likely than term-born infants to have a lower birthweight, a lower Apgar and higher Critical Risk Index for Babies II (CRIB-II) scores, receive postnatal antibiotics and be delivered by cesarean section (Table 1).

**Table 1.** Cohort Characteristics.

|  | Preterm | Term | P Value |
| --- | --- | --- | --- |
| Stooled < 48 h | 37 | 34 |  |
| Male Sex | 49% | 56% | 0.5580 |
| Black/African American | 78% | 91% | 0.1361 |
| Maternal Age (yr) | 26.94 ± 6.00 | 25.15 ± 4.64 | 0.1731 |
| Gestational Age (wk) | 27.51 ± 2.71 | 39 ± 1.15 | <0.0001 |
| Birthweight (g) | 874.1 ± 266.81 | 3115 ± 360.06 | <0.0001 |
| CRIB-II Score | 10 (8:13) | 0 (0:0) | <0.0001 |
| Apgar Score (5 min) | 6 (5:9) | 9 (9:9) | <0.0001 |
| Cesarean Delivery | 68% | 35% | 0.0058 |
| Prenatal Antibiotics | 32% | 38% | 0.5987 |
| Prenatal Antibiotic Doses* (n) | 1 (1:1.5) | 2 (1:2) | 0.2231 |
| Prenatal Antibiotic Doses (n) | 0 (0:1) | 0 (0:1) | 0.7943 |
| Postnatal Antibiotics | 92% | 0% | <0.0001 |
| 16S rRNA Positive | 92% | 94% | 0.5397 |
| ITS rDNA Positive | 88% | 65% | 0.0224 |

\*Number of doses received prenatally in those who received antibiotics. n (quartile 25%: quartile 75%) or n ± SD.

**Fig. 1.**
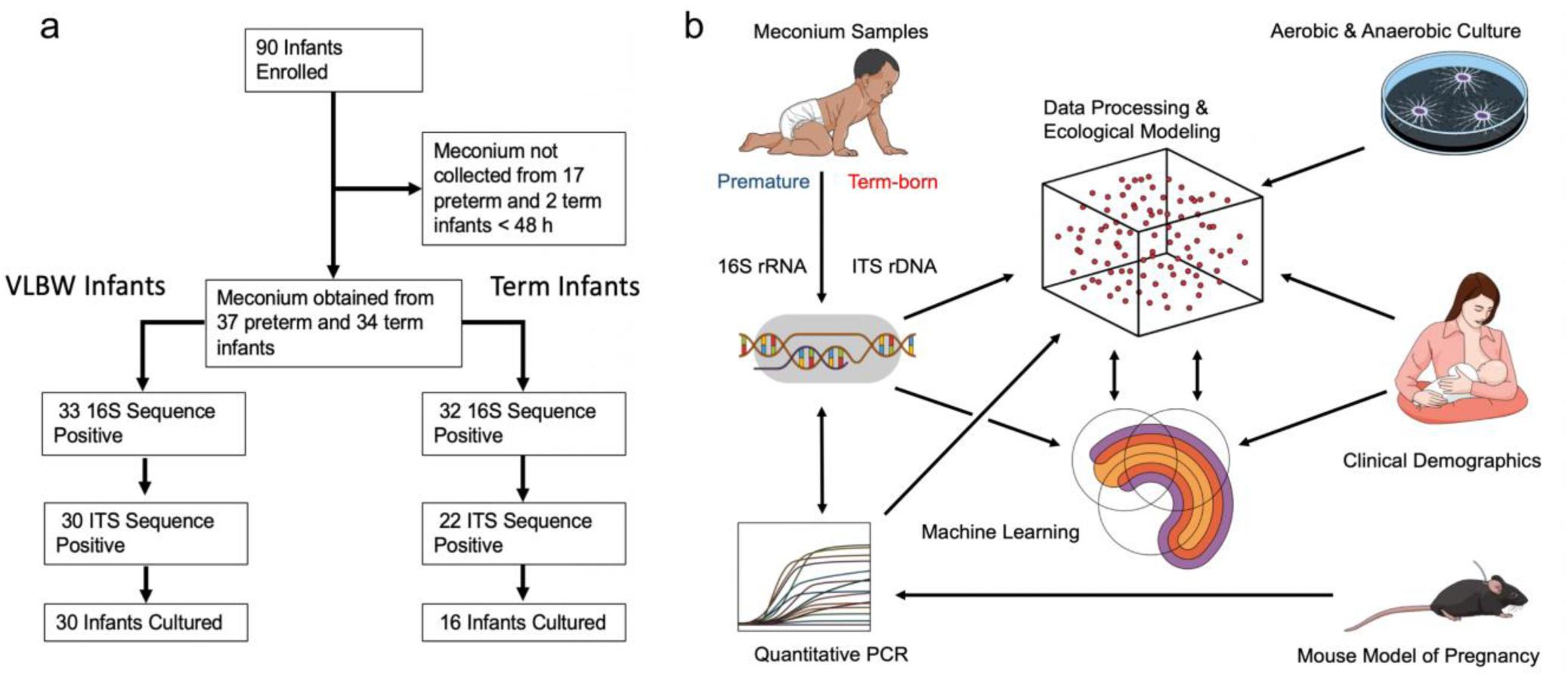
Curation of clinical and microbial data. **a** Schematic of patient allocation. VLBW, very low birthweight (preterm infants < 1500 g). **b** Schematic of data acquisition and analysis. Low biomass samples of meconium, an intralumenal material formed in the intestine prior to birth, were analyzed by culture-based techniques or Illumina MiSeq and quantitative PCR after high-fidelity DNA extraction. Sequence reads were subjected to exacting data processing to remove potential contaminants. Clinical demographics were used to perform ecological modeling of microbial communities and develop sequential rarified random forest classifier machine learning models. A mouse model of normal pregnancy was utilized to quantify intrauterine fungal biomass. 16S rRNA, 16S ribosomal RNA (bacteria and archaea). ITS rDNA, internal transcribed spacer ribosomal DNA (fungi).

### Fungal community composition differs by gestational age

The bacterial community composition of the microbiome differs by gestational age at birth^13^. This may be related to differences in initial meconium colonization^3^. To explore if fungal colonization of the microbiome was influenced by gestational age, we performed a binary comparison of infants born at <33 weeks’ (n = 37) and infants > 33 weeks’ gestation (n = 34). Meconium samples from preterm infants were more likely to contain fungal DNA than samples from term-born infants using internal transcribed spacer region 2 ribosomal DNA (ITS2 rDNA) amplicon sequencing at a read depth of 300x (88 versus 65%, χ^2^ *p* = 0.0224, Table 1). The alpha diversity of these fungal sequences clearly increased with increasing gestational age (Shannon diversity index, ANOVA *f* = 4.5 *p* =0.04. Principal coordinates analysis (PCoA) of Bray-Curtis dissimilarity matrices demonstrated fungal community structure was different by permutational multivariate analysis of variance (PERMANOVA, *R*^*2*^ =0.308 *p* =0.01, Fig. 2c). Discriminant analysis of principal components suggests that the principal variables at the genus level are the prevalence of *Elmerina, Cladosporium, Candida* and *Stereum*. Similarly, canonical correspondence analysis (CCA, χ^2^ distance = 0.71 *f* = 1.14 *p* =0.004), and redundancy analysis (RDA, variance =3.83 *f* =1.22 *p* =0.001) were both consistent with a clear differentiation in fungal community structure by advancing gestational age (Fig. 3d).

**Fig. 2.**
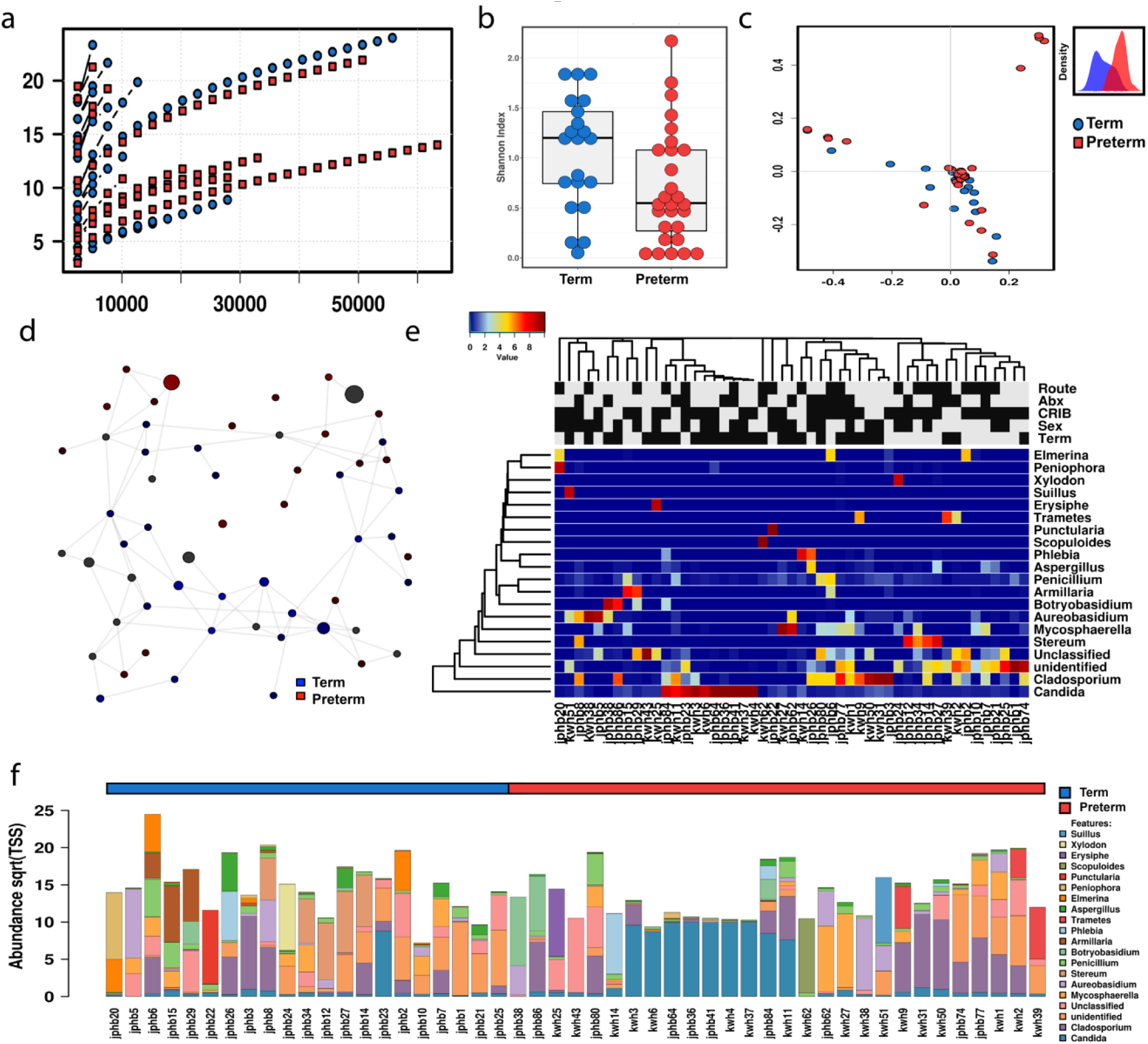
Fungal ecologies in human meconium vary by the gestational age at birth. **a** Sequencing rarefaction analysis of species richness. **b** Alpha diversity quantified by the Shannon index, ANOVA *f* = 4.5 *p* =0.04. **c** Principal coordinates analysis of Bray-Curtis dissimilarity matrices, PERMANOVA *R*^*2*^ =0.308 *p* =0.01. The subset displays a discriminant analysis of principal components. **d** Network analysis developed using Pearson’s correlations. Positive correlations with false discovery rate-adjusted, *p*-values <0.05 are presented as an edge. **e** Heat map of differential distribution of taxa at the order level, ranked by gestation. **f** Relative abundance of taxa at the phylum level. Sqrt(TSS), square root total sum normalization (Hellinger transformation). For all analyses n=71. Preterm samples are displayed in red and term-born samples in blue.

**Fig. 3.**
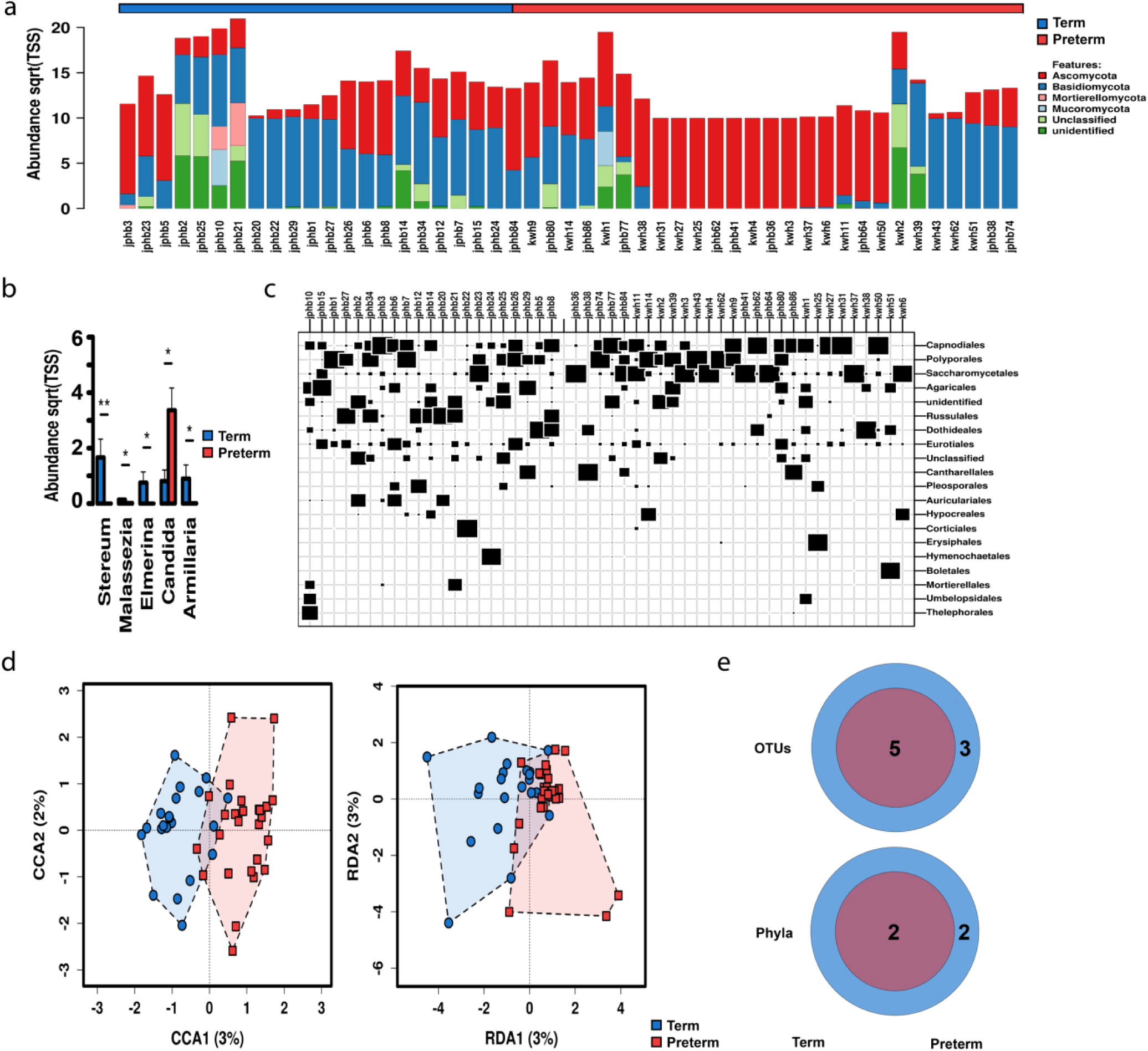
Fungal communities increase in complexity with advancing gestational age at birth. **a** Relative abundance at the phylum level. Sqrt(TSS), square root total sum normalization (Hellinger transformation). **b** Distribution of key taxa at the genus level. ANOVA, Bonferroni, * *p* < 0.05, \*\**p* < 0.01. **c** Twenty most abundant interkingdom taxa at the order level. **d** Canonical correspondence analysis (CCA), χ^2^ = 0.71 *f* = 1.14 *p* =0.004, and redundancy analysis (RDA), variance =3.83 *f* =1.22 *p* =0.001. **e** Core interkingdom OTUs and unique phyla between preterm and term-born infants. OTU, operational taxonomic unit. Data are median ± IQR (n=71). Preterm samples are displayed in red and term-born samples in blue.

Network analysis supports the sparseness of fungal communities, particularly in preterm infants, which is likely related to the low fungal diversity (Fig 2d). However, as quantified by positive Spearman’s correlations, fungal networks do appear to develop complexity with increasing gestational age.

The relative composition of the fungal communities varied with gestational age (Fig. 2 and 3). At the order level, preterm infants were primarily colonized with Saccharomycetales (73.4%), with other fungal taxa accounting for less than a quarter of assigned reads. Term-born infants possessed mycobiomes that were less dominated by a singles species than preterm infants. Saccharomycetales and Polyporales accounted still accounted for about half of the fungal burden in infants with a later gestation (22.1% and 27.7%, respectfully), but other minor taxa were more markedly prevalent with increasing gestational age. In addition to Polyporales (5.2% < 33 weeks’ and 22.1% > 33 weeks’ gestation), Russulales showed the most pronounced expansion with increasing gestational age (0.8% < 33 weeks’ and 14.5% > 33 weeks’ gestation, Fig. 2f, 3a). The most significant expansions in abundance at the genus level were in *Stereum, Malassezia, Elmerina* and *Armillaria*. In contrast to this trend, the domination of the mycobiome by *Candida* in some preterm infants lead to this particular genus being uniquely more abundant in preterm infants overall. Finally, core microbiome analysis also supports the expansion of fungal communities with increasing gestational age, with the addition of 2 unique core phyla and 3 unique core OTUs in term-born as compared to preterm infants (Fig. 3e).

To further investigate the association of specific fungal taxa with gestational age, we performed linear discriminant analysis of effect size (LEfSe) to identify fungal taxa that could perform as high-dimensional biomarkers for prematurity. The prevalence of the fungal order Saccharomyces and specifically the genus *Candida* reliably identified preterm samples. In addition, the prevalence of fungal class Elmerina and the genera *Stereum* and *Malassezia* reliably identified samples from full-term newborns (Supplemental Fig. 3i).

In childhood, fungal colonization has been shown to be influenced by host sex^14^. We examined fungal community composition in our cohort to see if fungal populations in meconium were influence by the sex of the host. Alpha diversity of was unaltered by sex (Supplemental Fig. 4g). Relative abundance differences were most pronounced for Polyporales (24.1% in females and only 6.9% in males), with other orders having less dramatic differences (Supplemental Fig. 4h). Meconium fungal community structure was not influenced by host sex (PCoA, PERMANOVA *R*^*2*^ =0.0209 *p* =0.529, Fig. 4a). Similarly, redundancy analysis (RDA, variance =3.19 *f* =1.02 *p* =0.348) was also not significant.

**Fig. 4.**
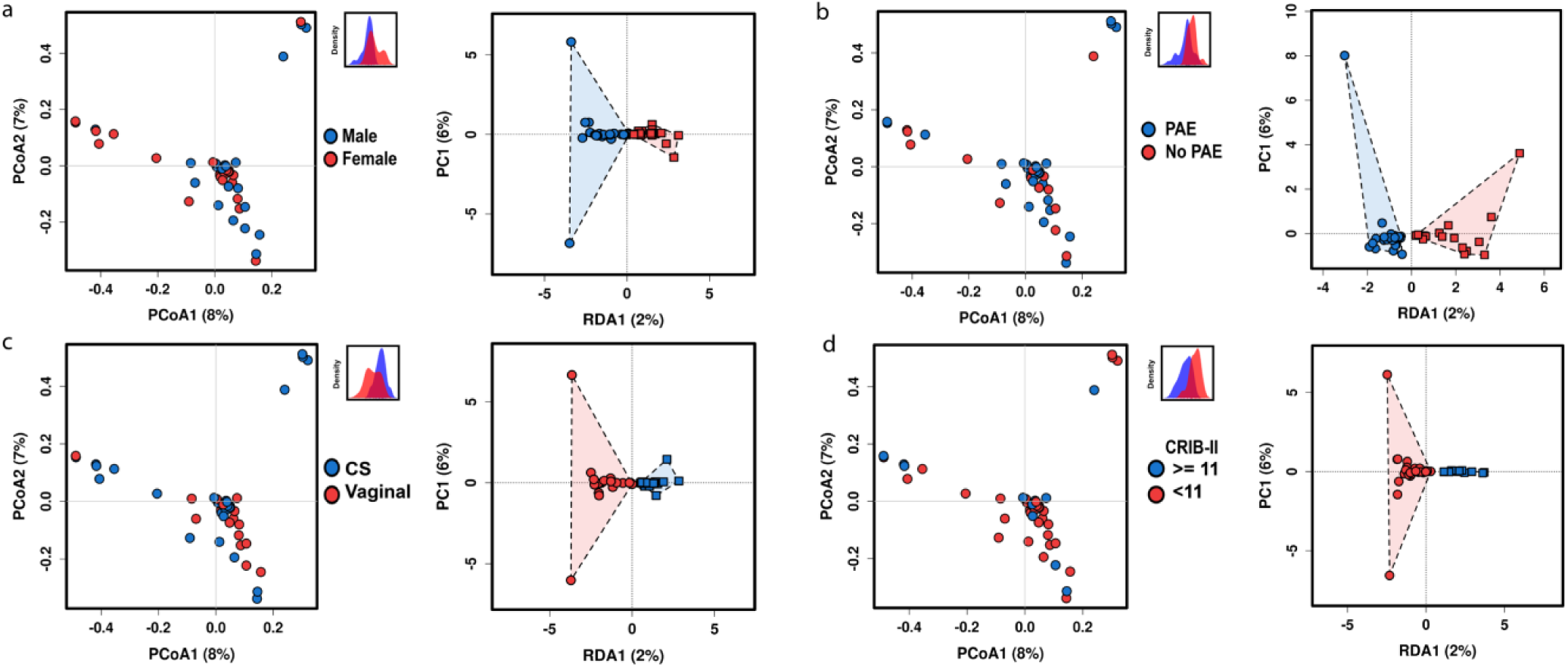
Fungal community structure is not determined by common perinatal factors. **a** Host sex does not significantly alter fungal community composition. Principal coordinates analysis (PCoA) of Bray-Curtis dissimilarity matrices, PERMANOVA *R*^*2*^ =0.0209 *p* =0.529. Redundancy analysis (RDA), variance =3.19 *f* =1.02 *p* =0.348. The subset displays a discriminant analysis of principal components. **b** Prenatal antibiotic exposure (PAE) does not significantly alter fungal community composition. PCoA of Bray-Curtis dissimilarity matrices, PERMANOVA *R*^*2*^ =0.202 *p* =0.598. RDA, variance =3.51 *f* =1.12 *p* =0.083. **c** Mode of delivery does not significantly alter fungal community composition. PCoA of Bray-Curtis dissimilarity matrices, PERMANOVA *R*^*2*^ =0.202 *p* =0.598. RDA, variance =3.51 *f* =1.12 *p* =0.083. **d** Illness severity does not significantly alter fungal community composition. PCoA of Bray-Curtis dissimilarity matrices, PERMANOVA *R*^*2*^ =0.0251 *p* =0.146. RDA, variance =2.68 *f* =0.85 *p* =0.947. CRIB-II, Clinical Risk Index for Babies II. For all analyses n = 71.

Antibiotics can have a profound effect on the compositional structure of the microbiome^15^, and lead to changes in the fungal colonization as well ^16^. To characterize the potential effect of maternal antibiotic exposure on the initial structure of the mycobiome, we examined how any antibiotic exposure altered fungal composition of the meconium. In our samples, perinatal antibiotic exposure did not produce significant differences in alpha diversity (Supplemental Fig. 4d). The relative abundance of Saccharomycetales was increased by maternal antibiotic exposure (60.1% after antibiotic exposure, and 32.5% without antibiotic exposure, Supplemental Fig. 4e). The distribution of other major taxa was similar. Fungal community structure was not significantly altered by prenatal antibiotic exposure (PERMANOVA *R*^*2*^ =0.202 *p* =0.598 and RDA, variance =3.51 *f* =1.12 *p* =0.083, Fig. 4b).

Whether mode of delivery influences microbial colonization remains controversial^2,17^. To explore influence of the delivery mode on fungal colonization we examined fungal community structure using cesarean or vaginal delivery as a binary variable. Alpha diversity was not significantly altered by mode of delivery, although cesarean born infants did demonstrate more variability (Supplemental Fig. 4a). Relative composition differences showed some variation by mode of delivery (Supplemental Fig. 4b) Similar to preterm infants, samples from infants born via caesarean were predominantly colonized by Saccharomycetales (61.7% after cesarean and 39.7% after vaginal). Both groups expressed similar levels of Polyporales (15.3% after cesarean and 13.8% after), but Russulales was more prevalent in vaginally-born infants (13.2% after vaginal and 0.62% after cesarean). While preterm infants were more likely to be delivered by cesarean in our cohort (68% versus 35% χ^2^ *p =*0.0058, Table 1), as characterized by PCoA, fungal community structure did not significantly differ after either vaginal or cesarean delivery (PERMANOVA *R*^*2*^ =0.202 *p* =0.598 and RDA, variance =3.51 *f* =1.12 *p* =0.083, Fig. 4c), which is consistent with the formation of meconium prior to birth.

Increasing illness severity closely associates with decreasing gestational age and is therefore a covariate and potential confounder in many diseases related to preterm birth^18^. To gauge a potential confounding influence of illness severity, we examined fungal community composition in relation to a critical score of 11 on the CRIB-II. No differences were appreciated in alpha diversity as quantified by the Chao1 index (Supplemental Fig. 4j). The relative abundance differences were also less markedly altered than for other perinatal factors (Supplemental Fig. 4k). The abundance of Saccharomycetales was unaltered at the class level, but the abundance of Polyporales was reduced (15.5% with score <11 and 6.9% with a score ≥11) and the relative abundance of Capnodiales and Dothideales increased with elevated CRIB-II score (6.0% <11 and 13.5%, ≥11; and 1.6% <11 and 10.8% ≥11, respectfully). Clinical illness severity did not appear to alter fungal community composition (PERMANOVA *R*^*2*^ =0.0251 *p* =0.146 and RDA, variance =2.68 *f* =0.85 *p* =0.947, Fig. 4d).

### Fungi form complex interkingdom communities

We examined how fungi formed interkingdom microbial communities by the characterizing the resulting community composition from the combined 16s and ITS sequencing results (n =71). For most infants, fungal communities were a minor component their interkingdom microbiome, but a few preterm infants had microbiomes that were dominated by fungi (Fig. 5 and Supplemental Fig. 1). This may emphasize the importance of pioneer species to the ecological succession of the primordial gut. In general, the interkingdom microbiome increased in complexity with increasing gestational age at birth. While rarefaction analysis favored more microbial richness in term samples (Fig. 5a), alpha diversity as quantified by the Shannon diversity index was not significantly altered (ANOVA *f* =2.2 *p* =0.14, Fig 5b). PCoA of the combined microbial communities showed clear differential clustering for with a binary cut off at 33 weeks’ gestation (Bray-Curtis dissimilarity matrices, PERMANOVA with 999 permutations *R*^*2*^ =0.103 *p* =0.000333, Fig. 5c). Discriminant analysis of principal components suggests that the principal variables at the genus level are the prevalence of *Methylobacterium, Bacteroides* and *Stereum*. The most prominent differences in abundance at the phylum level were in Blasidiomycota, Bacteroidetes and Actinobacteria, all of which were more abundant in term born infants. Acidobacteria, in contrast, were more prevalent in preterm infants (Supplemental Fig. 1b). At the genus level, there were expansions in multiple genera with increasing gestational age. The most prominent changes were in *Methylobacterium* and *Bacteroides* (Supplemental Fig. 1d). However, there were some genera, in particular *Lactobacillus* and *Staphylococcus*, that were more abundant in preterm infants, but at a much lower level. The expansion of the interkingdom microbiome with increasing gestation is also supported by analysis of the core microbiome of term infants which increased from 24 to 36 unique OTUs and added 7 unique phyla to the core preterm microbiome (Supplemental Fig. 1e).

**Fig. 5.**
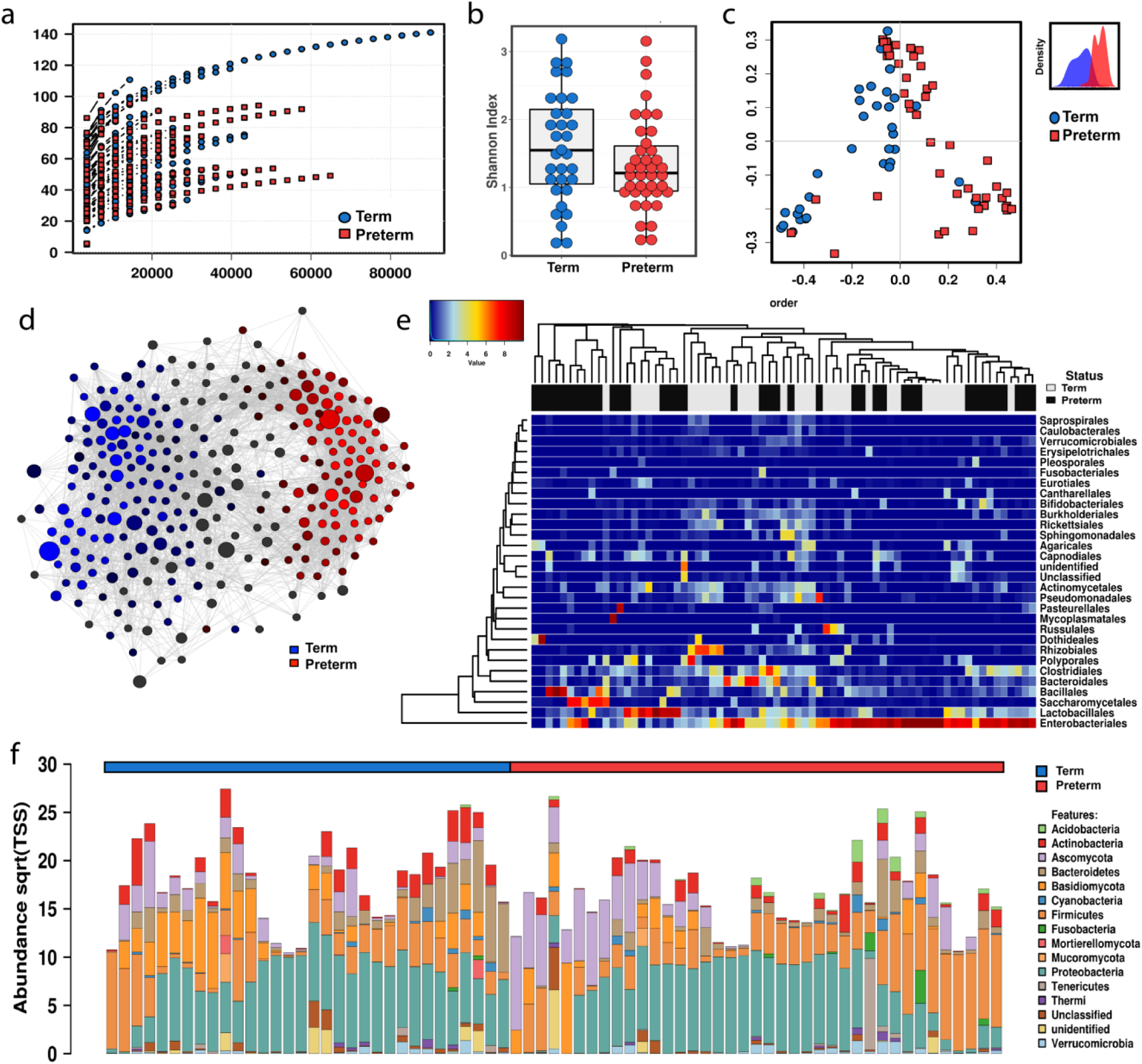
Fungi and bacteria form complex interkingdom microbial communities in human meconium. **a** Sequencing rarefaction analysis of species richness. **b** Alpha diversity quantified by the Shannon index, ANOVA *f* =2.2 *p* =0.14. **c** Principal coordinates analysis of Bray-Curtis dissimilarity matrices, PERMANOVA with 999 permutations *R*^*2*^ =0.103 *p* =0.000333. The subset displays a discriminant analysis of principal components. **d** Network analysis developed using Spearman’s correlations. Positive correlations with false discovery rate-adjusted, *p*-values <0.05 are presented as an edge. **e** Heat map of differential distribution of taxa at the order level, ranked by gestation. **f** Relative abundance of taxa at the phylum level. Sqrt(TSS), square root total sum normalization (Hellinger transformation). For all analyses (n=71). Preterm samples are displayed in red and term-born samples in blue.

**Fig. 6.**
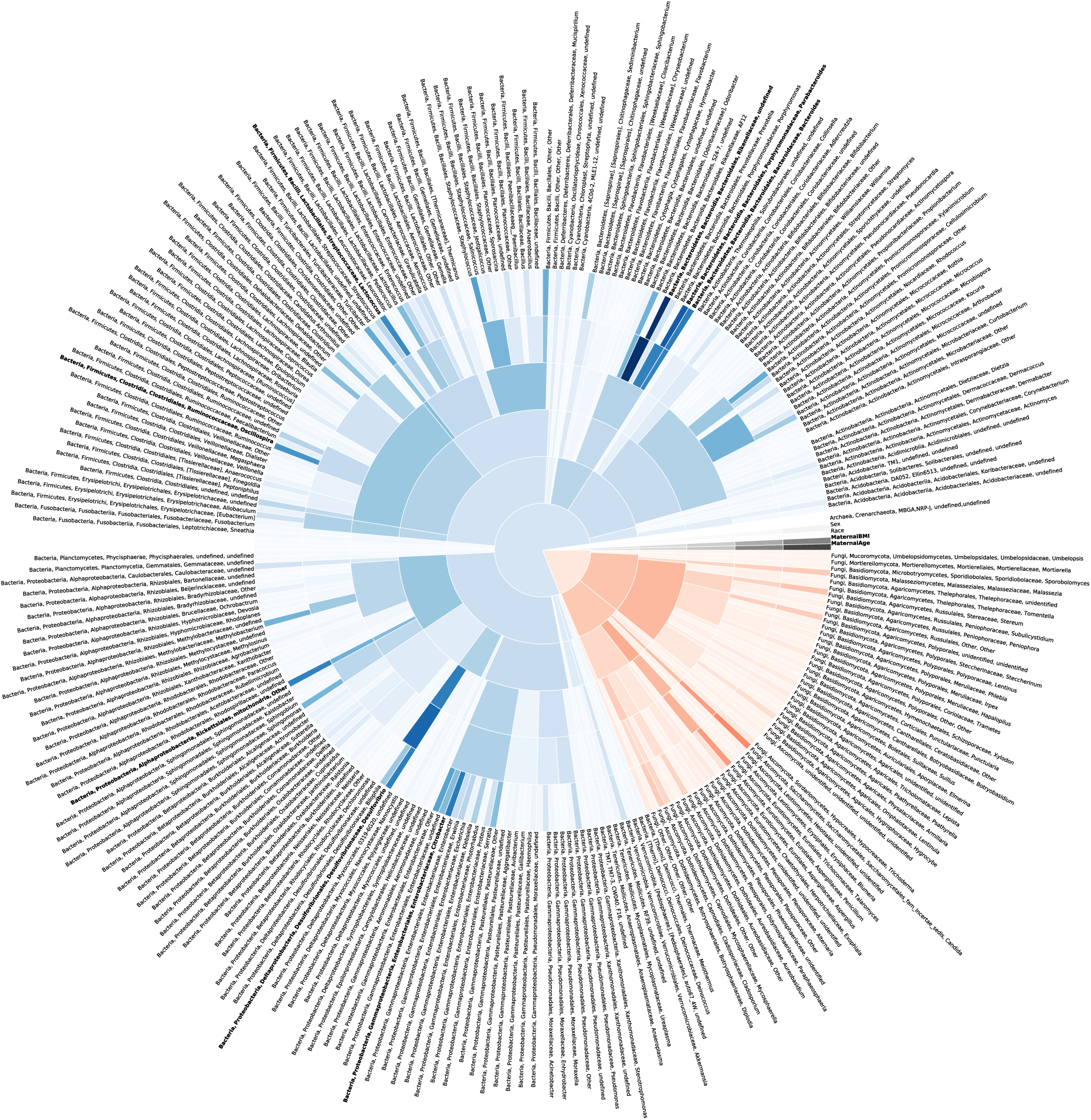
Interkingdom microbial community composition accurately classifies preterm infants. The precision of rarified random forest classifier machine learning models developed using microbial taxa from each level of the phylogenic tree increases as the phylogenic level decreases, highlighting the importance of key microbial taxa. Darker shading indicates a more important predictor to the machine learning model. Blue represents bacteria, red represents fungi and black represents archaea and clinical demographics. Starting from the center and moving outward, concentric rings represent models developed utilizing taxa on the kingdom, phylum, class, order, family, and genus levels. The 10 most important predictors based on the genus-level model are highlighted in bold. Area under the curve statistics for each level are reported in Supplementary Fig. 6b.

To understand interrelatedness in interkingdom ecology, we used network analysis. Compared to the sparse inter-fungal networks, fungi and bacteria form complex interkingdom networks with numerous interactions (Figure 5d). The expanding networks led to multiple unique nodes clearly separated by gestational age at birth.

To characterize the influence of perinatal factors on interkingdom community structure, we used PCoA and RDA to capture the impact of host sex, prenatal antibiotic exposure, mode of delivery and illness severity. Similar to fungal community structure, the interkingdom structure was not significantly altered by these common perinatal exposures as assessed by both constrained and unconstrained analyses (Supplemental Fig. 2).

### Key interkingdom community members predict gestational age of the host

To explore the distribution of microbial taxa with respect to gestational age at birth, we developed a machine learning model to predict the gestational age of a newborn based on the interkingdom composition of the microbiome. We developed our first machine learning model using only microbial taxa. Starting with 51 fungal and 209 bacterial taxa that were identified at least twice in the dataset, we utilized LASSO with 5-fold cross validation to identify potential significant predictors, leading to the identification of 9 fungal and 34 bacterial taxa clearly distributed by gestational age (Supplemental Fig. 6a). A support vector machine model reliably classified infants as preterm or term utilizing these taxa (Supplemental Fig. 6b).

### Interkingdom community composition classifies preterm status

To understand the relative strength of key microbial taxa to function as predictors of preterm status relative to other clinical demographic data, we implemented a random forest model with 5-fold cross validation using the presence of 260 genus-level microbial taxa and select clinical demographics. After 100 independent permutations, average sensitivity analysis of the final random forest model provided an area under the curve (AUC) of 0.859 [95% confidence interval (CI) 0.857-0.862, Supplemental Fig. 6c]. We then performed a variable importance analysis on the random forest model and selected the top 10 variables to build a rarified random forest model using 5-fold cross validation. After 100 permutations, the average performance of the sensitivity analysis of the rarified random forest models had an AUC 0.867 (95% CI 0.861-0.871). Overall, the strongest predictors from these key taxa ran counter to the general trend and were more abundant in preterm than in term-born infants (Undefined genera within family Rikenellaceae, and the bacterial genera *Parabacteroides* and *Citrobacter*). With the exception of *Lactococcus*, the other key genera were more abundant in term-born infants. Of note, maternal body mass index and age also functioned as reliable predictors, which may be related to their known association with preterm birth^19^.

We compared the random forest models to a variety of other classifiers. All other methods compared provided classification accuracy < 71%, which was significantly lower than the accuracy of the random forest classifiers (Supplemental Fig. 6d). During model development, we did not include clinical variables such as prenatal steroid use, duration of rupture of membranes, antibiotic use and delivery mode as predictors because such variables are closely associated with impending delivery and would confound predictive modeling. However, we implemented Mann-Whitney-U (or Kruskal-Wallis) tests to determine whether there were significant differences between the presence of these variables and the 10 key microbial taxa we previously identified during development of the random forest model. We found no significant differences between antibiotics use, duration of membrane rupture or delivery mode and presence of these 10 taxa. In contrast, there were statistically significant differences (*p* <0.01) between use of prenatal steroids (none, 1 or 2 doses) and the abundance of 7 of these key taxa (Supplemental Fig. 7a).

To quantify the performance of higher-order microbial taxa, we developed models exploring the strength of different phylogenic levels to function as predictors of prematurity. We designed our modeling approach in six levels to capture the influence of using increasingly more granular taxa information on the classification accuracy. The accuracy of the model increased as the as the phylogenic level of the taxa used decreased, highlighting the importance of the distribution of key tax and their variation with gestational age (Fig. 5 and Supplemental Fig. 7b). In addition, when using higher level microbial taxa as potential predictors, the mothers’ age and body mass index were relatively more important as predictors. These statistical models demonstrate the differential variation of the interkingdom community composition and underscores the close relationship between the microbial community composition and gestational age of the host at birth.

### Fungal and microbial organisms can be cultured from the primordial microbiome

To explore if our next generation sequencing-based analyses were potentially representative of live microorganisms as opposed to translocated microbial DNA, we performed anaerobic and aerobic culture-based assays for both fungi and bacteria on a subset of samples (n = 41) for which sufficient meconium was available (Supplemental Fig. 8). Aerobic fungi-selective culture media was positive in 32% of preterm and 29% of term infant samples (χ^2^ *p* = 0.835). Aerobic bacteria-selective culture media was positive in 89% of preterm and 71% of term infant samples (χ^2^ *p* = 0.128). Anaerobic fungi-selective culture media was positive in 50% of preterm and 21% of term-born infant samples (χ^2^ *p* = 0.079). Anaerobic bacteria-selective culture media was positive in 69% of preterm and 88% of term infant samples (χ^2^ *p* = 0.1543). Overall, bacterial cultures were diffusely positive, while fungal cultures were less likely to demonstrate growth. While contamination cannot be completely excluded, we cannot eliminate the possibility of live fungi and bacteria in the primordial human gut.

### The fetal gut contains fungal DNA

Considerable controversy persists as to whether the isolation of microbial DNA from the human placenta or newborn gut is representative of live microbial organisms^20^. While the similarities between microbes observed in the placenta and neonatal meconium suggest transfer of microbial DNA across the placenta, they do not represent direct observation of the fetal intestine. However, bacterial DNA has been identified in the fetal gut in a mouse model of normal pregnancy^21^. To directly investigate the fetal intestine for the presence of fungal DNA and live organisms we developed a mouse model of normal pregnancy. Using C57BL6J mice, we collected samples from the fetal gut and several maternal body sites on either estimated gestational day 17 (E17) or within 24 h of birth (P1). We used real time PCR to identify ITS biomass across all maternal and fetal samples. The relative fungal biomass was about 14-fold lower in the fetal intestine before birth than in the maternal amniotic fluid. However, fungal biomass was about 6-fold more abundant in the neonatal intestine than the amniotic fluid (mean cycle threshold, C_t_ = 36.75 ±15.48 at E17 and C_t_ = 25.71 ±15.16 at P1 and C_t_ = 29.20 ±6.58 in amniotic fluid).

Dectin-1 is a pattern recognition receptor that recognizes the fungal cell-wall β-glucan and is important for detection of fungal pathogens in humans and mice^22^. To investigate if inhibited host sensing of fungi altered the transplacental deposition of fungal biomass in the neonatal gut, we collected samples from the fetal gut and multiple maternal body sites from dectin-1^-/-^ mice on the C57Bl6J background (B6.clec7a^tm1Gdb^). In the fetal intestine, dectin-1^-/-^ mice had about a 6-fold increase in fungal biomass over dectin-1 competent mice (C_t_ = 33.50 ± 14.81 versus C_t_ = 36.75 ±15.16). However, after birth, the neonatal intestinal fungal biomass was lower on P1 (C_t_ = 29.33 ±18.01 versus C_t_ = 25.71 ±15.48) and remained a more significant proportion of the intestinal biomass. While there are considerable differences between the placentas of humans and mice^23^, these results suggest that the intrauterine transfer of fungal DNA may be widespread across mammals. Impaired host-sensing of fungi may alter both intrauterine deposition and post-natal colonization.

## Discussion

Here we present the first description of the primordial fungal ecology of the neonatal gut using both next-generation sequencing and culture-based techniques. Microeukaryotes, primarily fungi, were present in the first-pass meconium samples collected within hours of birth in both preterm and term-born newborns. In general, as quantified by amplicon sequencing of eukaryotic ITS rDNA, infantile fungal communities developed greater taxonomic diversity as the gestational age at birth increased. The structure of these communities was unaffected by perinatal factors, such as mode of delivery, antibiotic exposure, infant sex, and illness severity, consistent with intrauterine formation of meconium. Culture-based techniques indicate microbial DNA may originate from viable microorganisms present in the neonatal gut.

Understanding the primary succession of microorganisms into the human gut is important because the order in which these potential colonizers are introduced into the intestine influences the mature community structure. Colonization experiments in mice suggest the timing of arrival of microbial colonizers is the major factor in determining final community structure^24^; which helps explain why other perinatal factors, such as perinatal antibiotic exposure, account for only limited variability in mature community structure. Bacterial communities then undergo a rapid expansion during the first several years of life^25^, eventually establishing a host-specific core microbiome that resists further perturbations^26^. Because misconfiguration of the core microbiome during this period likely contributes to multiple host disease states, understanding the ecological processes driving microbiome assemblage is crucial to developing therapeutic interventions^27^. Recent evidence suggests that the mycobiome remains relatively constant during the first year of life^11,12^, which underscores that the primary succession of initial fungal colonizers may be even more important for final community structure than the succession of bacterial species. In addition, fungi may also form a crucial nidus for bacterial colonization^6^. Our evidence for the presence of commensal fungi in the primordial gut and the development of these communities with gestational age demonstrates this process may begin earlier that was previously thought - with implications that extend throughout life.

Meconium forms during early human gut development, eventually migrating into the colon around the 19^th^ week of gestation^28^. Passage of meconium is a developmentally programmed event that usually occurs within 48 hours after birth^28^. Once enteral feeding has been established environmental microbes invade the gastrointestinal tract and are detectable in the first true stools. Until the development of next-generation sequencing techniques, the presence of microorganisms in the first-pass meconium was opaque using traditional culture-based techniques. Meconium, and therefore the uterine environment, has been widely accepted to be sterile^9^. However, during the past decade several researchers have demonstrated bacterial DNA can be isolated from meconium^3,8,29-34^ and the placenta^35-37^, but if this microbial biomass is representative of live organisms remains controversial. These studies have focused on the bacterial components of meconium, however, leaving fungi undescribed.

The human amniotic fluid may also harbor a very low level of microbial organisms. Chu et al.^17^ have demonstrated that the bacterial components of the meconium microbiome are distinct and do not resemble adult vaginal and skin communities like the predominant bacterial members of other neonatal body sites. Our results show interkingdom microbial communities that are essentially unaffected by perinatal exposures, with the possible exception of prenatal steroids. The demonstration of live microorganisms in the newborn gut represents a monumental shift from the sterile womb hypothesis that dominated perinatal biology for the last century^8,9^. The increasing prevalence and the development of bacterial ecology with increasing gestational age suggests that this is a natural, non-pathogenic phenomenon^3,33,34^. The human fetus regularly swallows and replenishes the amniotic fluid throughout gestation, which suggests a mechanism for how rare microbes in the amniotic fluid could become more concentrated within the primordial gut. Based on these findings, we postulate that low abundant amniotic fluid bacterial and fungal biomass is continuously collected during gestation and accumulates in the intestine. By the time of delivery, meconium may contain a collective record of fetal microbial exposure in an analogous manner to how meconium contains a record of transplacental illicit drug exposure^38^. This process would also expose the fetal mucosal immune system to low levels of microbial products that may initiate host microbial surveillance mechanisms required to process the influx of microbes into the gut that occurs during and after delivery.

To directly observe the fetal environment, we utilized a mouse model of normal pregnancy. This work highlights the presence of fungal biomass within the developing fetal gut. The implication of a steady deposition of microbial biomass in the fetal intestine suggests that this is a natural process, which raises the question of whether translocation of microbes is regulated. To gain insight into this process, we asked if impairment of the host’s recognition of fungi would alter the deposition of fungal biomass within the fetal and neonatal gut. Indeed, we found that loss of the host fungal pattern recognition receptor Dectin-1 resulted in more fungal biomass within the fetal intestine. While there are marked differences between the placentas of humans and mice^21^, this suggests that the deposition of fungal biomass within the fetal gut may be a regulated process that requires host recognition of fungal organisms.

Counter to the general trend of our results which support an apparently natural process of microbial biomass accumulation in the neonatal intestine, our finding that some preterm infants are dominated by fungal order Saccharomyces, and in particular the genus *Candida*, may support the alternate hypothesis that in some instances preterm birth is a product of polymicrobial disease of the uterine cavity^39^. After birth, systemic infection with *Candida* are a persistent threat during neonatal care. *Candida* species, primarily *Candida albicans*, are opportunistic pathogens particularly associated with bacterial dysbiosis^40^. *Candida* are significant commensals of the human gut^41^, so the presence of *Candia* within the gastrointestinal tract is understandable. In the mature gut, commensalism is maintained by a tripartite interaction between *Candida*, other members of the microbiome, and host immunity^41^. However, *Candida* are also the dominate fungal species associated with microbial invasion of the amniotic cavity^42^, and are detectable in the amniotic fluid by culture and culture-independent techniques^43^. As quantified by culture-based techniques, *Candida* chorioamnionitis is a rare but accepted cause of preterm birth^44^. Our study showed that preterm meconium samples were more likely to contain fungi than term infants. Furthermore, the mycobiomes of preterm infants were also more likely to be dominated by *Candida*, as opposed to the more diverse mycobiomes of infants with a longer gestation. Regardless of these concerning findings, conclusively demonstrating that *Candida* colonization is associated with preterm birth will require a clinical trial powered to test this hypothesis.

Fungi are ubiquitous in the environment. In low biomass samples like meconium, the presences of environmental fungi are difficult to separate from organisms originating within the samples. Because microeukaryotic DNA is a relatively low proportion of the fecal DNA content, especially in meconium, exacting extraction, purification and amplification techniques are required. A prominent reason that investigation of the mycobiome has lagged behind investigation of the microbiome is a lack of standardization in detection methods for fungal organisms^41^. Our choice to utilize amplification of the ITS2 region, the read depth and the database used to align our amplicon reads, therefore, could all influence the fungal species identified in our samples. Incidentally, the exacting DNA extraction techniques we utilized may also explain the differences in bacterial community structure with prior work because they increased the detectability of low biomass samples. We froze meconium samples immediately after collection to prevent post-excretion microbial expansion that would alter the relative composition of microbial communities, and passed negative control samples throughout the sample collection, DNA extraction and amplicon sequencing process. However, the fact that perinatal exposures known to shape the post-natal microbiome did not significantly alter community composition in our samples argues that significant post-natal microbial colonization did not occur before sample collection.

Our findings support the presence of fungal organisms within the neonatal gut. Taken together, our sequencing and culture-based techniques support the presence of live fungal organisms in meconium in addition to bacteria. Homeostasis of fungal commensals may represent an important aspect of human biology present even before birth with functional implications throughout early life. Low abundance colonization of the primordial gut with interkingdom ecologies appears to be a normal part of healthy pregnancy that might represent a priming of the intestine for the exponential expansion of the microbiome after birth. The domination of microbial communities by fungal order Saccromycetes in some preterm infants, however, suggests a potential pathologic association with preterm birth.

## Methods

### Study patients

Written and informed consent was obtained from the infants’ mothers. All investigations were carried out in accordance with the declaration of Helsinki under The University of Tennessee Health Science Center Institutional Review Board protocol 17-05311-XP. We prospectively enrolled either very low birthweight infants (< 1500 g) preterm or term-born (estimated gestational age 37-41 weeks) infants (Fig. 1a). We excluded any mother-infant dyads with the potential for immune deficiency, but otherwise invited all mothers that meet our entrance criteria during the study period to participate.

### Infant exposures

Data was collected regarding infant demographics and the following clinical exposures: prenatal and postnatal antibiotic exposure, perinatal steroid exposure, delivery mode, and illness severity (the CRIB-II^18^). Additional clinical characteristics were also abstracted from the medical record to characterize potential co-variates with gestational age.

### Sample collection

The first-pass meconium was collected during the first 48 h after birth. Samples were collected from diapers using sterile spatulas, placed into sterile microcentrifuge tubes and frozen at −30C within minutes of collection. Several times a week, batches of samples were collected and frozen at −80C until further analysis could be performed. Infants that did not stool during the first 48 h or for which a meconium sample was not collected were excluded from downstream analysis.

### DNA extraction and Illumina MiSeq sequencing

Meconium samples were resuspended in 500 μL of TNES buffer containing 200 units of lyticase and 100 μL of 0.1/0.5 (50/50 Vol.) zirconia beads. Incubation was performed for 20 min at 37°C. Following mechanical disruption using ultra-high-speed bead beating, 20 μg of proteinase K was added to all samples, and they were incubated overnight at 55C with agitation. Total DNA was extracted using chloroform isoamyl alcohol, and total DNA concentration per mg stool was determined by qRT-PCR. Purified DNA samples were sent to the Argonne National Laboratory (Lemont, IL) for amplicon sequencing using the NextGen Illumina MiSeq platform. Blank samples passed through the entire collection, extraction and amplification process remained free of DNA amplification.

### Bioinformatics

Sequencing data were processed and analyzed using QIIME (Quantitative Insights into Microbial Ecology) 1.9.1. Sequences were first demultiplexed, then denoised and clustered into sequence variants. For bacteria we rarified to a depth of 3,000 sequences. (In combined analysis after excluding samples with a bacterial depth of < 3,000, we used a depth of 300 for further analysis).

Representative bacterial sequences were aligned via PyNAST, taxonomy assigned using the RDP Classifier. Processed data was then imported into Calypso 8.84 for further analysis and data visualization^45^. The Shannon Index was used to quantify alpha diversity (inter-sample)^46,47^. Unweighted UniFrac analysis was used to quantify beta diversity (intra-sample) ^48^, and the differences were compared using PERMANOVA with 999 permutations. For fungi, sequences were aligned, and taxonomy was assigned using the UNITE (dynamic setting) database^49^. Fungal OTUs were rarified at a depth of 300 sequences for alpha diversity using Shannon or Chao1 and beta diversity using Bray-Curtis dissimilarity or Jaccard abundance ^50^ indices. As with bacteria, beta diversity significance was then assessed using PERMANOVA. To quantify relative abundance of taxa between groups, we utilized ANOVA adjusted using the Bonferroni correction and FDR for multiple comparisons. We used LEfSe to test for significance and perform high-dimensional biomarker identification^51^. Network analysis was generated from Spearman’s correlations. Positive correlations with an FDR-adjusted, *p* <0.05 were presented as an edge.

### Machine learning and variable selection

In the first phase, we developed a support vector machine utilizing 34 bacterial and 9 fungal taxa identified utilizing an unbiassed LASSO approach with 5-fold cross validation and generated a confusion matrix to characterize the performance of the model as a predictor of gestational age

In the second phase, we developed a random forest-based classification model to distinguish between preterm and term-born newborns using maternal age, maternal body mass index, newborn sex and 260 microbial taxa that were present in at least two samples in the data set. Random forest classifiers are based on the arrangement of multiple decision trees built using a random sample of predictors. The outcome of a random forest is based on a weighted average of the outcomes of individual decision trees within the model. To identify and control for potential overfitting, we implemented a 5-fold cross validation strategy. By repeating this process 5 times for each subject, we obtained a single class prediction that is purely based on a classifier built on a distinct dataset. We then repeated the development of the entire model 100 times. We also implemented a recursive variable selection based on the out-of-bag predictor importance analysis. Here, variable importance was assessed by randomly permitting values of a single predictor and quantifying the increment in the out-of-bag error rate. Variables that caused no drop on the error rate were assumed to have no influence in predicting outcome. We then selected the 10 most important variables and built another 100 permutations of our random forest model using 5-fold cross validation.

We also compared the performance of the random forest classifier we generated using this method to other classifiers including logistic regression, discriminant analysis, k-nearest neighborhood, decision trees, support vector machines and neural networks using AUC statistics. Finally, we calculated commonly used classification accuracy statistics such as specificity, sensitivity, and positive predictive value for the final random forest model to characterize the ability of the model to distinguish between preterm and term-born babies.

In the final phase, we repeated this model development process for each phylogenic level to understand the variable importance of different taxa. To achieve this, in level 1 (kingdom) we used four demographic variables and 3 microbial kingdoms (bacteria, fungi, and archaea) features by aggregating all bacterial and fungal taxa under their particular kingdom definition. Therefore, at level 1 (kingdom), we built a random forest classifier using a total of 6 features. The four demographics variables included host sex (female: 0, male: 1), maternal self-reported race/ethnicity (Black: 0, other: 1), maternal age, and maternal BMI. Fungal and bacterial taxa were represented in six different levels following, kingdom > phylum > class > order > family > genus. For level 2 (phylum) we aggregated taxa data at 23 distinct phylum-level taxa resulting in total of 27 features and including the same four demographics data. We repeated this process until we had reached the 6^th^ level (genus) where we utilized a total of 265 variables (4 demographics and 261 taxa data at genus level). At each level, we included only taxa that were observed in at least two different subjects to avoid overfitting. For each level, we implemented five-fold cross validation. To avoid sampling bias, we repeated the cross-validation after randomly shuffling the data for 100 times at each level. Classification performance was summarized as AUC statistics.

### Microbial culture

We prepared culture broth using either brain heart infusion (BHI) with 250 mg/L of fluconazole (bacteria-selective) or yeast-extract-peptone-dextrose (YPD) with 50 mg/mL of chloramphenicol (fungal-selective) under both aerobic and anaerobic conditions. Meconium samples were thawed and diluted in 1 mL of PBS. Tubes containing 1 mL of culture broth were inoculated with 100 μL of room temperature meconium diluted in 1 mL of culture broth. Samples were then incubated at 32°C under either aerobic or anaerobic conditions. Bacterial samples were assessed by optical density after 48 h and fungal samples after 120 h.

### Statistical analysis

We compared demographic and clinical variables between infants with a gestational age < 33 weeks’ and > 33 weeks’ using the χ^2^ or two-tailed Welch *t*-test for parametric variables or the Wilcoxon rank-sum test for non-parametric variables, as appropriate.

### Animal model of pregnancy

All animal experiments were approved by the University of Tennessee Health Science Center Institutional Animal Care and Use Committee. Timed-mating of adult C57BL6J mice or dectin-1^-/-^ on the C57Bl6J background (B6.clec7a^tm1Gdb^, Jackson Labs stock #012337) at 8-10 weeks of age was performed to obtain offspring samples at embryonic day 17 (E17) or post-partum day 1 (P1). After euthanasia, swabs were collected from the maternal mouth, skin and genitourinary tract. The dam was then soaked in exspor (chlorine dioxide disinfectant). Gravid uteri and intestines were removed from prepped dams in a UV-light sterilized laminar flow hood by sterile gowned and gloved personnel extensively trained in sterile technique (a maternal-fetal medicine specialist gynecologist and a neonatologist) using sterile technique and instruments that were sterilized between animals. Utilizing a second set of sterilized instruments, pups were removed, euthanized and placed in exspor then passed to another operator who removed them in a second laminar flow hood. The body cavity was opened with another set of sterile instruments and removed the intestine *en bloc* and placed them into a sterile container. Samples of the placenta, amniotic fluid, small intestine and colonic contents were also placed in sterile containers. All samples were snap frozen and stored at −80C until DNA extraction.

## Supporting information

Combined Supplemental Data File

## Data Availability

Microbial sequence data is available at: [To be uploaded on Revision]

## Computer Code

Custom computer code is available upon request.

## Author Contributions

KW, AT and JFP conceived of the study design; KW and JHP recruited the patients and collected the meconium samples; KW and EM extracted clinical data from the medical record; CG and JFP performed the DNA purification; KW, BP and JFP performed the culture-based analyses; KW and MA performed the murine experiments, IK and OA performed the statistical modeling and developed the machine learning models; KW, IK, OA and JFP performed the data quality control, statistical analysis and prepared the figures; KW wrote the paper; KW, MA, JHP, OA, BP, AT and JFP edited the manuscript; and all authors approved the final version.

## Acknowledgements

The authors would like to thank the mothers and infants who contributed to the study and to our research coordinators, Gail Camp and Nancy Ruch, for their assistance in identifying and recruiting patients.

