## Supplementary material for "Fungi form interkingdom microbial communities in the primordial human gut that develop with gestational age": Combined Supplemental Data File

Willis et al.

### Supplemental Text 1

**Bacterial colonization of the meconium**

The bacterial colonization of first-pass meconium has been previously described[3,20,21]. Unlike previous work, both preterm and term-born newborns were equally likely to contain bacterial DNA on 16S rRNA sequencing at a read depth of 3000x (90% <33 weeks' and 94% >33 weeks' gestation, $\chi^2$ *p* =0.5397, Supplemental Fig. 5b). Gestational age also did not alter the alpha diversity of meconium samples (Supplemental Fig. 5e). The most marked alteration in the relative composition of bacteria at the order level was in Bacteroidia (1.3% < 33 weeks' and 10.6% > 33 weeks' gestation) and Alphaproteobacteria (0.08% <33 weeks and 8.3% >33 weeks' gestation). In general, with increasing gestational age the relative abundance of rarer bacterial orders increased, with the exception of order Bacilli which decreased from 26.9% in preterm infants to 17.6% in those born at term (Supplemental Fig. 5f). Gestational age significantly altered the bacterial community structure (PERMANOVA of unweighted UniFrac distances with 999 permutations, *f*-statistic 4.65, *p* =0.001, Supplemental Fig. 5h).

We also performed LeFSe to identify bacterial taxa that could function as high-dimensional biomarkers of gestational age. The abundance of multiple bacterial genera identified preterm samples, including *Dermacoccus*, *Parabacterodies*, *Clostridium*, *Oribacterium*, *Anaerococcus*, *Citrobacter*, *Enterobacter* and *Erwinia*. The abundance of two taxa identified full term samples: family *Micrococcacea* and genus *Methylobacterium* (Supplemental Fig. 5j).

The alpha diversity was not significantly altered by any perinatal factor analyzed (Supplemental Fig. 7). As with fungi, mode of delivery was not associated with changes in bacterial community structure (PERMANOVA, *f*-statistic 1.192, *p* =0.19, Supplemental Fig. 7a). After antibiotic exposure, the relative abundance of order Gammaproteobacteria increased (46.8% without perinatal antibiotics exposure and 64.7% with exposure), while the relative abundance of order Bacilli decreased (27.8% without exposure and 14.4% with exposure, Supplemental Fig. 7e). However, overall bacterial community structures were not significantly altered by perinatal antibiotic exposure (PERMANOVA, *f*-statistic 1.47, *p* =0.052, Supplemental Fig. 7f). Far less prominent changes in relative abundance were noted by host sex (Supplemental Fig. 7h), and no differences were appreciated in bacterial community structure (PERMANOVA, *f*-statistic 1.18, *p* =0.179, Supplemental Fig. 7I). Finally, illness severity as quantified by the CRIB-II score was also not associated with significant differences in bacterial community structure (PERMANOVA, *f*-statistic 1.49, *p* =0.051, Supplemental Fig. 7l).

### Supplementary Figure 1

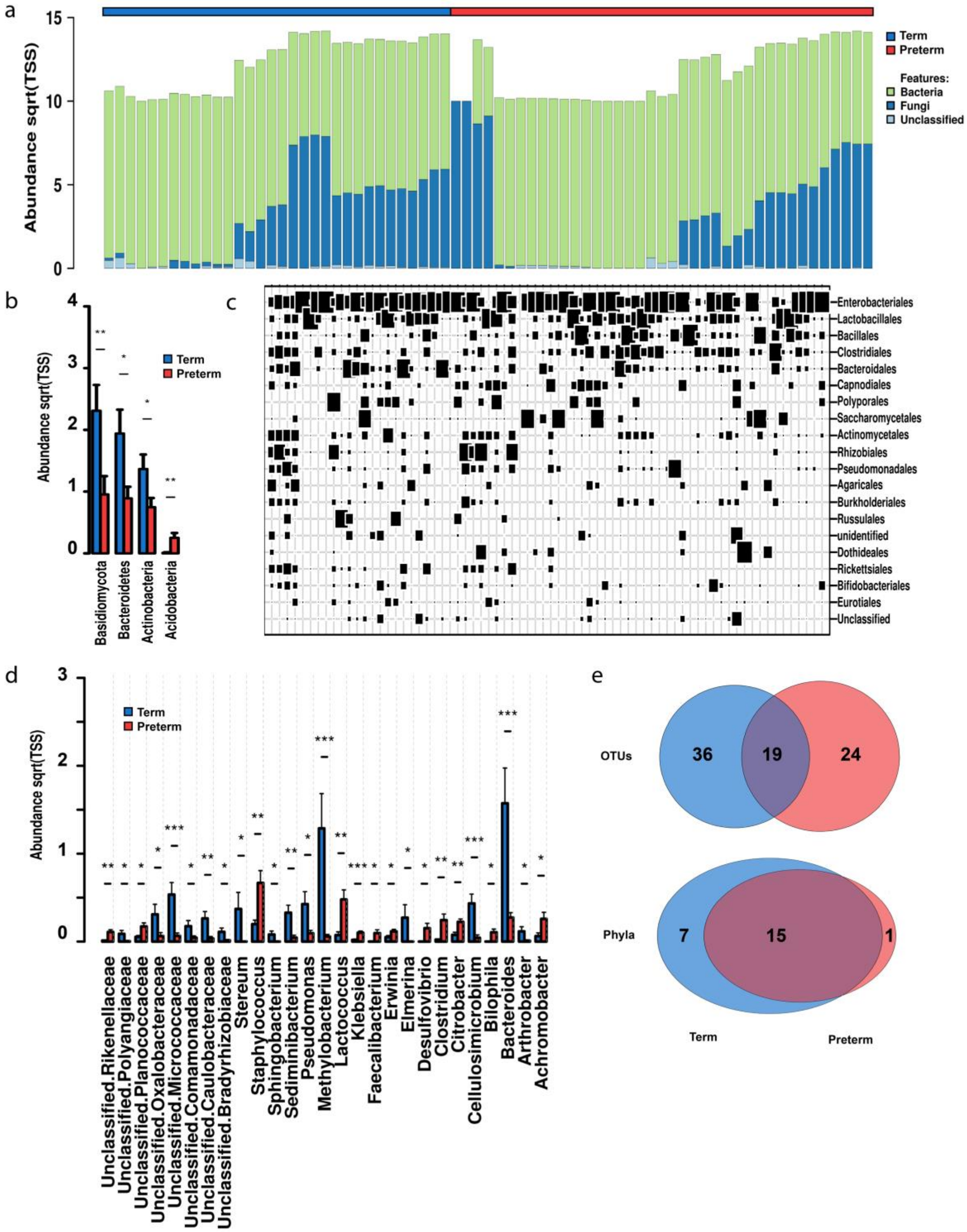


**Supplemental Fig 1.** Fungal and bacterial communities increase in complexity with advancing gestational age at birth.

**a** Superkingdom relative abundance. Bacteria are displayed in green and fungi are displayed in blue.

**b** Distribution of key taxa at the phylum level. Sqrt(TSS), square root total sum normalization (Hellinger transformation).

**c** Twenty most abundant interkingdom taxa at the order level.

**d** Distribution of key taxa at the genus level.

**e** Core interkingdom OTUs and unique phyla between preterm and term-born infants.

Data are median $\pm$ IQR (n=71). Preterm samples are displayed in red and term-born samples in blue.

For both (**a, d**) ANOVA, Bonferroni, * $p < 0.05$, ** $p < 0.01$, *** $p < 0.001$.

### Supplementary Figure 2

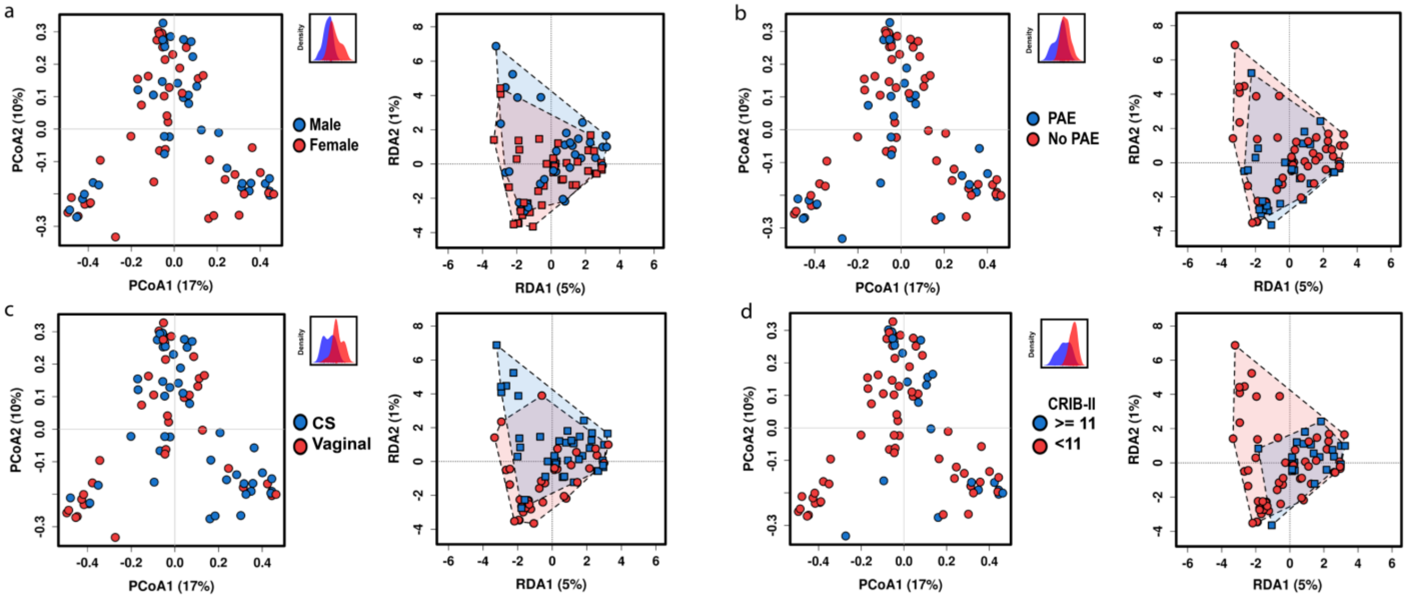


**Supplemental Fig. 2.** Interkingdom community structure is not determined by mode of delivery, prenatal antibiotic exposure, host sex or illness severity.

**a** Principal coordinates analysis (PCoA) of Bray-Curtis dissimilarity matrices showing host sex does not significantly alter community structure, PERMANOVA $R^2$ =0.00977 $p$ =0.749. Redundancy analysis (RDA), variance =4.21 $f$ =0.90 $p$ =0.710. The subset displays a discriminant analysis of principal components.

**b** Prenatal antibiotic exposure (PAE) does not significantly alter community structure, PCoA of Bray-Curtis dissimilarity matrices PERMANOVA $R^2$ =0.0101 $p$ =0.699. RDA variance =4.24 $f$ =0.91 $p$ =0.691.

**c** Mode of delivery does not significantly alter community structure, PCoA of Bray-Curtis dissimilarity matrices PERMANOVA $R^2$ =0.0216 $p$ =0.0526. RDA variance =5.34 $f$ =1.14 $p$ =0.169.

**d** Clinical illness severity does not significantly alter community structure, PCoA of Bray-Curtis dissimilarity matrices PERMANOVA $R^2$ =0.00718 $p$ =0.964. RDA variance =3.16 $f$ =0.68 $p$ =0.997. CRIB-II, Critical Risk Index for Babies II.

Gestational age (data not shown), RDA variance =17.14 $f$ =3.67 $p$ =0.001.

For all analyses n = 71.

### Supplementary Figure 3

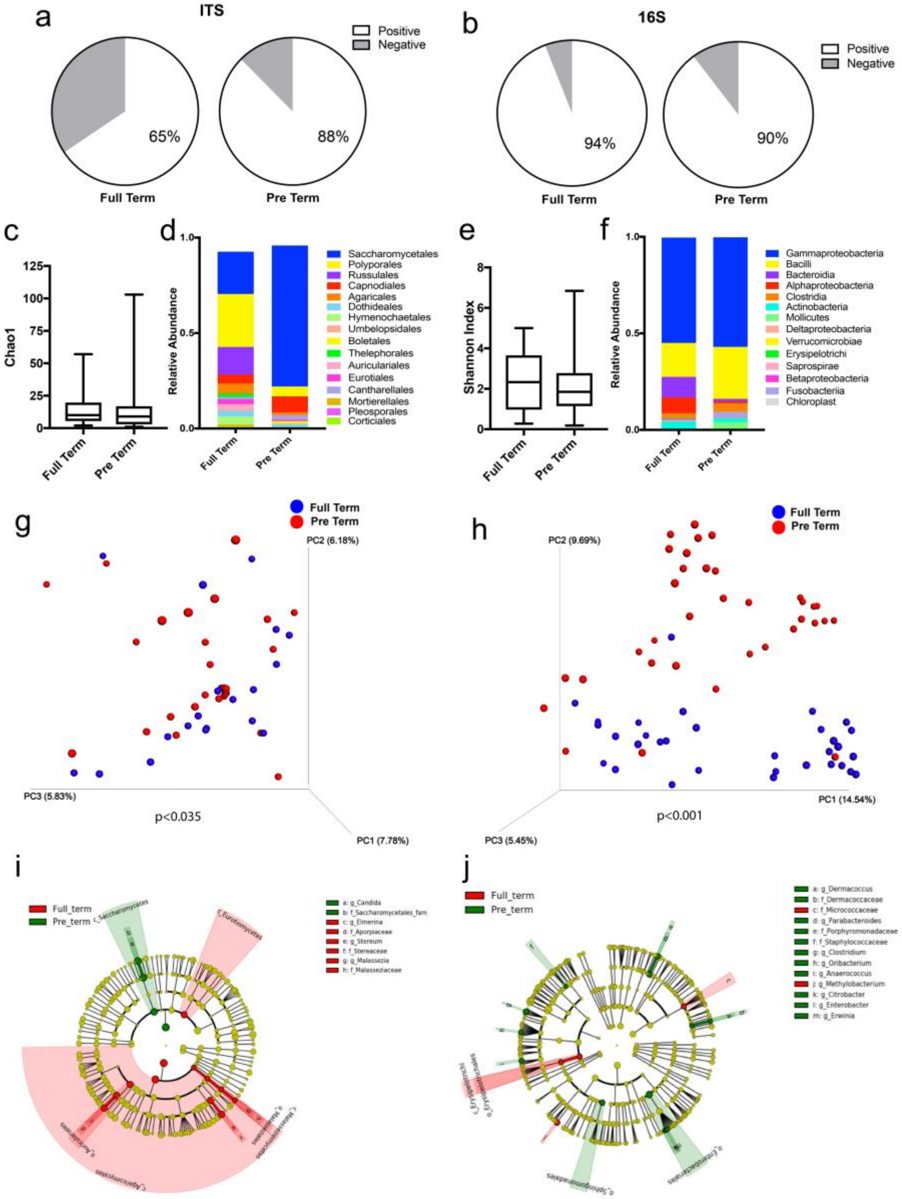


**Supplementary Fig. 3.** QIIME analyses showing bacteria and fungi form complex interkingdom ecologies.
**a** Percentage positive for ITS rDNA (fungi).
**b** Percentage of samples positive for 16S rRNA (bacteria and archaea).
**c** Alpha diversity quantified by Chao1 Index.
**d** Fungal class level taxonomy.
**e** Alpha diversity quantified by the Shannon Index.
**f** Bacterial class level taxonomy
**g** Principal component analysis of Jaccard distances, permutational multivariate analysis of variance (PERMANOVA) $p < 0.035$.
**h** Principal component analysis of unweighted UniFrac distances, PERMANOVA $p < 0.001$.
**i** Cladogram display of significantly enriched fungal taxa identified through linear discriminate analysis of effect size by gestational age.
**j** Cladogram display of significantly enriched bacteria taxa identified through linear discriminate analysis of effect size by gestational age.
Term-born infant samples displayed in red while preterm are displayed in green.
Data are median ± IQR (n=71).

### Supplementary Figure 4

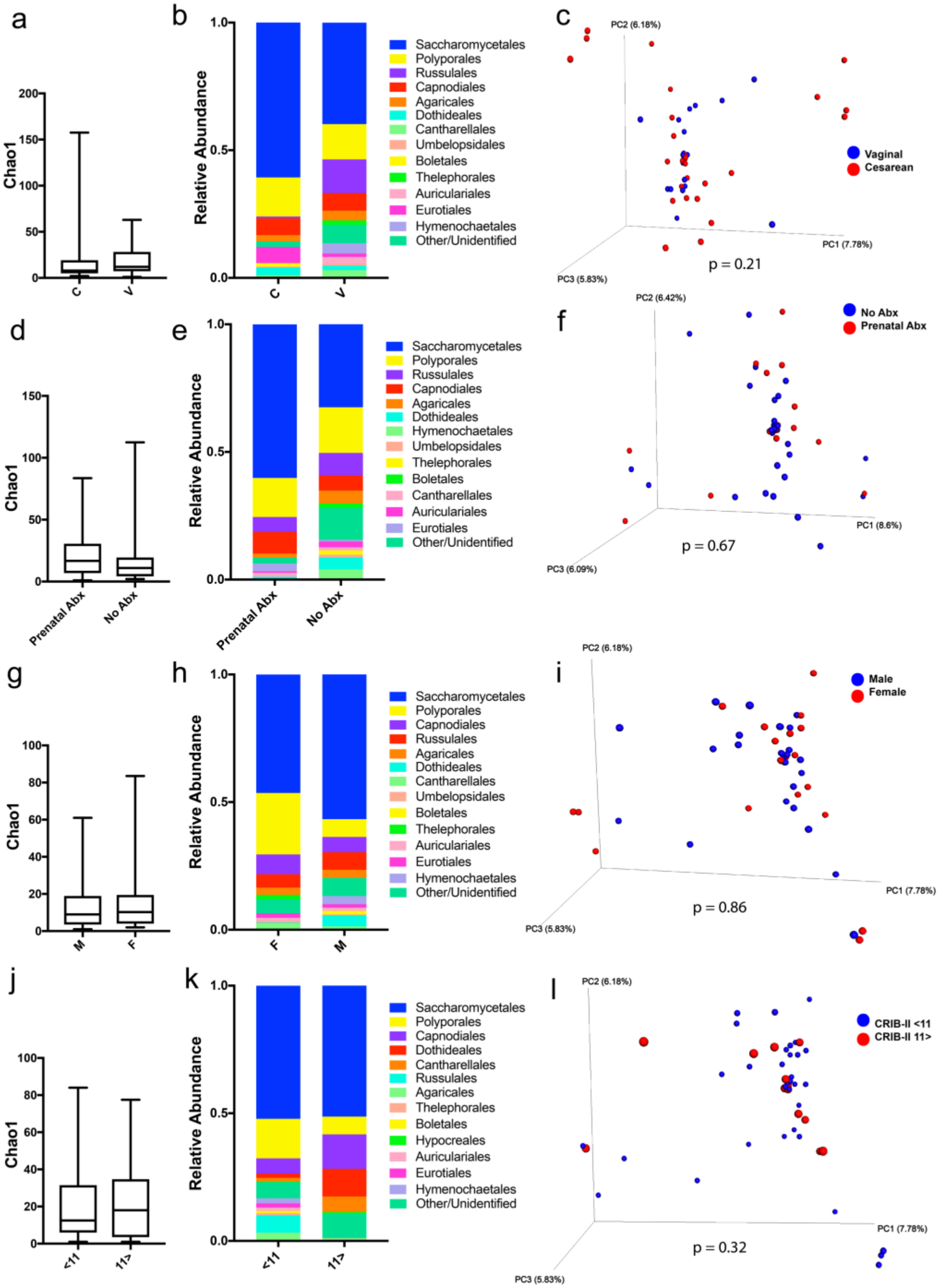


**Supplementary Fig. 4.** QIIME analyses showing fungal community structure is not determined by mode of delivery, prenatal antibiotic exposure, host gender or illness severity.

Chao1 indices of alpha diversity based on (**a**) on mode of delivery (MOD), (**d**) prenatal antibiotic exposure (PAE), (**g**) host gender and (**j**) illness severity as quantified by the Critical Risk Index for Babies II (CRIB-II).

Taxonomic relative distribution of bacterial communities based on (**b**) MOD, (**f**) PAE, (**h**) host gender, (**k**) and illness severity.

Principal component analysis of Jaccard abundance indices of fungal communities based on (**c**) MOD, with significance determined by permutational multivariate analysis of variance (PERMANOVA) $p$ =0.21; (**l**) PAE, PERMANOVA $p$ =0.67; (**l**) host gender, PERMANOVA $p$ =0.86; and (**l**) illness severity, PERMANOVA $p$ =0.32.

Data are median ± IQR (n=71).

### Supplementary Figure 5

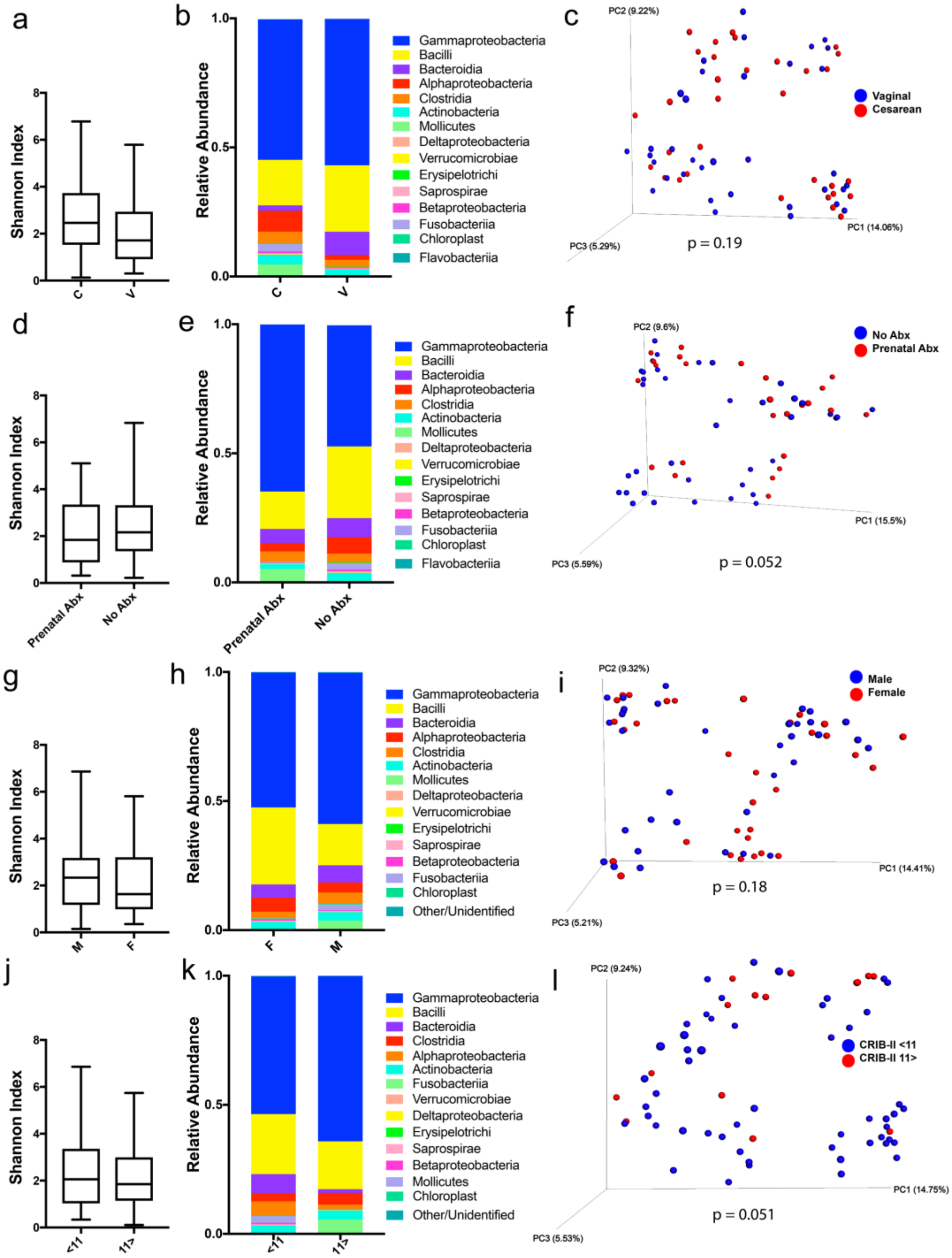


**Supplemental Fig. 5.** QIIME analyses showing bacterial community structure is not determined by mode of delivery, prenatal antibiotic exposure, host gender or illness severity.

Shannon indices of alpha diversity based on (**a**) on mode of delivery (MOD), (**d**) prenatal antibiotic exposure (PAE), (**g**) host gender and (**j**) illness severity as quantified by the Critical Risk Index for Babies II (CRIB-II).

Taxonomic relative distribution of bacterial communities based on (**b**) MOD, (**e**) PAE, (**h**) host gender, (**k**) and illness severity.

Principal component analysis of unweighted UniFrac distances of bacterial communities based on (**c**) MOD, with significance determined by permutational multivariate analysis of variance (PERMANOVA) $p$ =0.19; (**f**) PAE, PERMANOVA $p$ =0.052; (**i**) host gender, PERMANOVA $p$ =0.179; and (**l**) illness severity, PERMANOVA $p$ =0.051.

Data are median ± IQR (n=71).

### Supplementary Figure 6

a

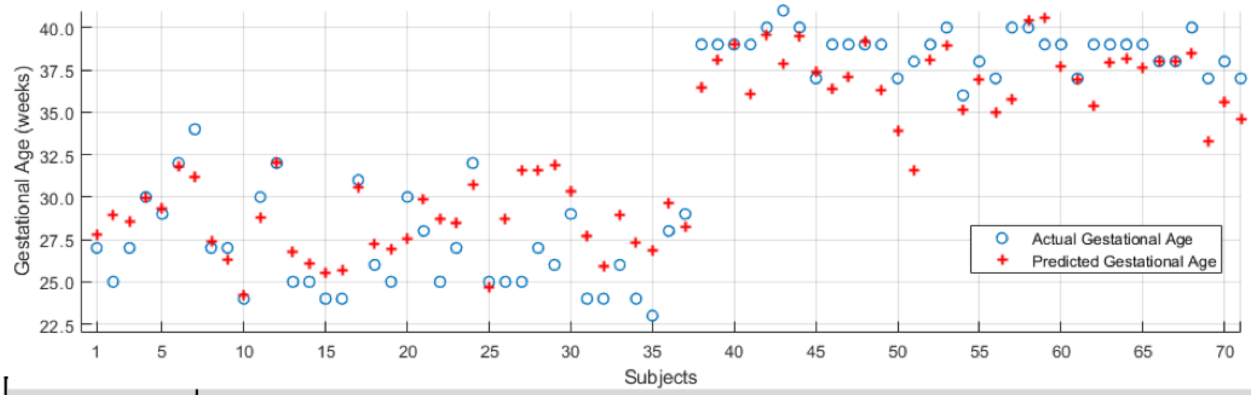


b

| Model 1 | Predicted | | | |
|---|---|---|---|---|
| | **Category** | **Term** | **Preterm** | **Accuracy (%)** |
| **Actual** | **Term** | 31 | 6 | Specificity = 83.8 |
| | **Preterm** | 2 | 32 | Sensitivity = 94.1 |
| | **Predictive Value (%)** | NPV = 93.9 | PPV = 84.2 | Overall = 88.7 |

c

| Actual / Predicted | Term | Preterm | Accuracies |
|---|---|---|---|
| **Term** | 29.0 | 5.0 | Specificity = 85.3% |
| **Preterm** | 5.1 | 31.9 | Sensitivity = 86.2% |
| | Negative Predicted Value = 85.1% | Positive Predicted Value = 86.5% | Accuracy = 85.8% |

d

| Algorithm | AUC with 96% CI (5-fold cross-validation results) |
|---|---|
| **Decision Trees** | 0.70 (0.58, 0.82) |
| **Logistic Regression** | 0.44 (0.31, 0.57) |
| **Support Vector Machines** | 0.65 (0.52, 0.78) |
| **K-nearest neighbors** | 0.56 (0.43, 0.69) |

**Supplemental Fig. 6.** Machine learning models to predict gestational age using microbial taxa.
**a** Utilizing 51 fungal and 209 bacterial taxa that were identified at least twice in the dataset, we utilized LASSO with 5-fold cross validation to identify significant predictors, leading to the identification of 9 fungal and 34 bacterial taxa as predictors of gestational age yielding an $R^2$ of 0.85 and an adjusted $R^2$ of 0.62.
**b** Confusion matrix from a support vector machine model utilizing these key microbial taxa.
**c** Confusion matrix from the final genera-level random forest machine learning model.
**d** Comparison of other machine learning models with less accuracy than random forest.

### Supplementary Figure 7

**a**

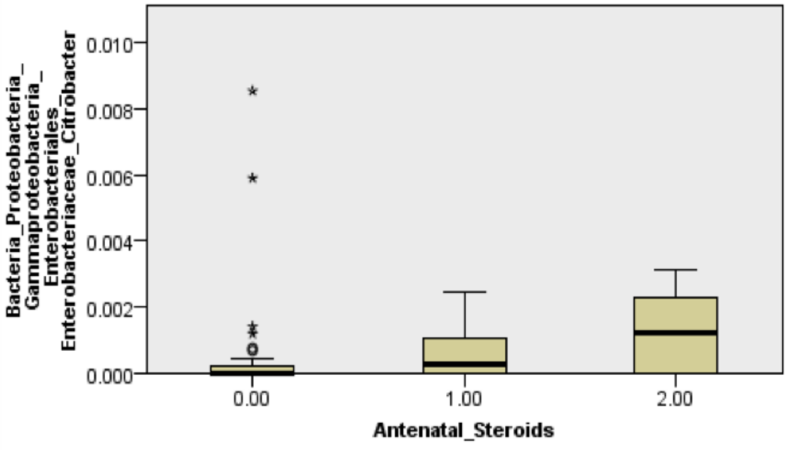


**b**

| Analysis Level | Accuracy (95% CI) | Number of Features |
|---|---|---|
| Level 1: Kingdom level | 57.63% [56.81%-58.44%] | 7 |
| Level 2: Phylum Level | 68.22% [67.41%-69.02%] | 27 |
| Level 3: Class Level | 74.27% [73.56%-74.98%] | 53 |
| Level 4: Order Level | 79.04% [78.43%-79.66%] | 87 |
| Level 5: Family Level | 84.22% [83.65%-84.79%] | 167 |
| Level 6: Genus Level | 85.74% [85.23%-86.24%] | 265 |

**Supplemental Fig. 7.** Machine learning models of interkingdom community importance.
**a** Box plot showing the alteration of 7 of the 10 most important bacterial taxa by the use of antenatal steroids as quantified by independent sample Kruskal-Wallis tests.
**b** Confusion matrix from a random forest classifier model for each phylogenic level.

### Supplementary Figure 8

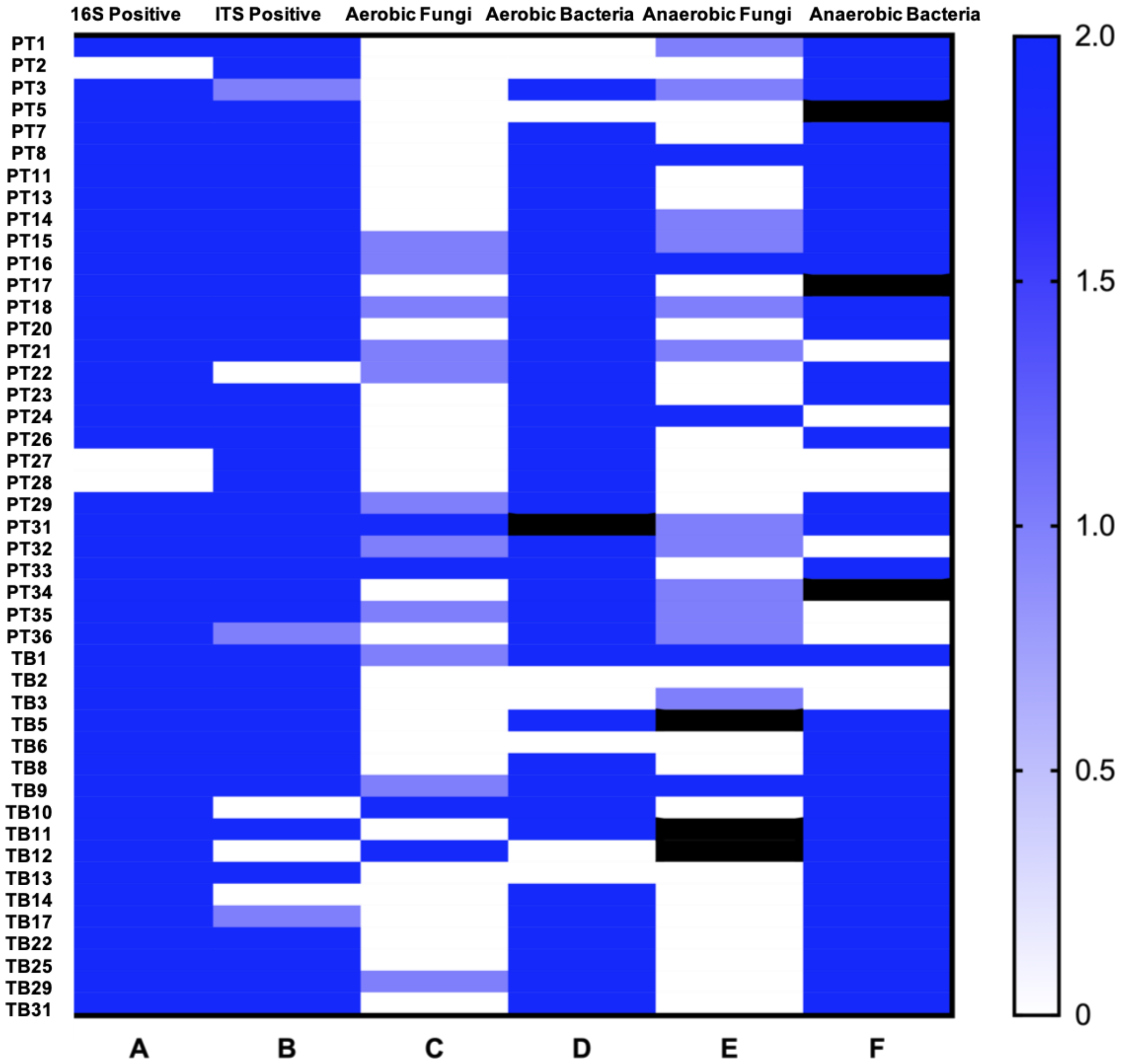


**Supplemental Figure 8.** Live fungi and bacteria are present by culture-based techniques.
**a** 16S rRNA (bacteria and archaea).
**b** ITS rDNA (fungi).
**c** Aerobic yeast-extract-peptone-dextrose (YPD) with chloramphenicol broth (aerobic fungi).
**d** Aerobic brain heart infusion (BHI) with fluconazole broth (aerobic bacteria).
**e** Anaerobic YPD with chloramphenicol broth (anaerobic fungi).
**f** Anaerobic BHI with fluconazole (anaerobic bacteria).
2 = strongly positive, 1 = weakly positive, 0 = undetected. Black cells represent quantity not sufficient.
(n = 46; PT, preterm n = 30; TB, term-born n = 17)
